# Likelihood Ratios for physical traits in forensic investigations

**DOI:** 10.1101/2024.05.25.595720

**Authors:** Franco Marsico, Thore Egeland

## Abstract

Recent years have seen significant advances in DNA phenotyping, which predicts the physical traits of an unknown person, such as hair, eyes, and skin color, using DNA data. This technique is increasingly used in forensic investigations to identify missing persons, disaster victims, and suspects of crimes. A key contribution of DNA phenotyping is that it allows researchers to search through lists of individuals with similar characteristics, often gathered from testimonies, photographs, and social media data. However, despite their growing relevance, current methods lack comprehensive mathematical models to calculate likelihood ratios that accurately assess the statistical weight of evidence. Our work bridges this gap by developing new likelihood ratio models, validated through computational simulations. In addition, we demonstrate the ability of these models to improve forensic investigations in real-world scenarios. Furthermore, we introduce the R package `forensicolors`, freely available on CRAN, to facilitate the application of the methodologies developed.

## 1 Introduction

In forensic investigations, the search for missing persons remains a critical and complex challenge. Salado Puerto et al. [1] introduced key definitions for this field. A Missing Person (MP) is defined as an individual with known identity but whose body remains missing. In contrast, an Unidentified Person (UP) is either a living individual with unknown biological identity, a common problem related to child abduction [2], or an unidentified human remains. The search process involves collecting both genetic and non-genetic data. For MPs, data include testimonies, legal documents, family interviews, social media, dental and medical records. For UPs, data are derived from unidentified remains and the context of their discovery, such as cemetery records with burial and death dates. More recently, DNA analysis has been expanded to infer various phenotypic characteristics, including skin, eye and hair color, as well as age [3, 4, 5, 6].

The studies by Evett and Weir [7, 8] established fundamental principles for interpreting forensic evidence. They highlighted the essential role of forensic scientists in determining probabilities of evidence based on various hypotheses. Central to this approach is the application of Bayes Theorem, a structured method for updating initial beliefs with new evidence to arrive at a posterior probability of the hypotheses in light of the data. In forensic investigations such as missing person and disaster victim searches, the comparison of two hypotheses is typical. The Likelihood Ratio (LR), defined as the ratio of the probabilities of observing the data under each hypothesis, quantifies the strength of evidence in favor of one hypothesis over another [7]. This method is particularly relevant in the statistical evaluation of DNA evidence in missing person cases, where the common contrasting hypotheses are: *H*_1_, the Unidentified Person (UP) is the Missing Person (MP); and *H*_2_, UP and MP are unrelated [9]. In the search for missing persons, DNA samples can be obtained directly from the UPs, such as human remains or individuals with uncertain biological identities. In the case of MPs, where direct samples are often unavailable, DNA from relatives is utilized and kinship testing is carried out [10]. This DNA analysis requires accurate knowledge of the family relationship between the MP and their relatives, often involving genotyping more than one relative. These relationships are typically depicted in family pedigrees, a crucial tool in genetics [11, 12].

Genetic databases have increasingly become a fundamental resource in kinship analyzes involving multiple pedigrees and UPs [2]. The field of large-scale DNA-based identification, including forensic applications such as Familial Search, Missing Person Identifications (MPI), Disaster Victim Identifications (DVI) and criminal investigations, have witnessed substantial advancements. This progress was fueled by technological innovations in software and the development of new genetic markers [13, 14, 15, 16]. MPIs typically represent large-scale operations with an indeterminate number of missing individuals, thus classified as open cases. In contrast, DVIs generally involve a known number of victims, which is categorized as closed cases [9]. Forensic experts face various complexities in both DVI and MPI scenarios, including the analysis of samples from charred remains, the grappling with scarce preliminary data, and the analysis through DNA database searches characterized by low statistical power [1, 10, 17].

In particular, for closed cases, Vigeland et al. showed that identifying each UP individually can give inconsistent solutions and proposed a methodology that jointly considers all UPs [18]. In all these cases, increasing the statistical power in hypothesis testing is imperative, which could require the incorporation of additional family members or genetic markers [2, 10, 19]. Also, the integration of non-genetic information can facilitate the prioritization of certain cases for intensified genetic scrutiny and hypothesis formulation [20, 21, 22, 23].

The inclusion of non-genetic information (such as age, gender, pigmentation traits, among others) in identifying missing persons is crucial. For example, it enables researchers to search through lists of individuals with similar characteristics, often gathered from testimonies, photographs, and social media data. However, its formal mathematical integration has been relatively unexplored [14, 20, 21, 23]. Recent developments have introduced models to calculate likelihood ratios for different types of non-genetic data [23]. Kinship analysis tools, such as Familias [24], use non-genetic factors to help filter unidentified persons based on missing persons characteristics. With regard to physical traits, research points to the importance of considering geographical differences to enhance the accuracy of DNA predictions [4]. This approach improves the search process and helps form more accurate hypotheses, especially when direct DNA samples are not available. Advances in (epi) genetics, particularly in the estimation of age, have added another layer of precision to these investigations [6]. Techniques such as DNA methylation analysis could provide valuable insights that help refine the search for missing individuals [25]. However, it is important to correctly statistically weigh these new methods and not to rely only on filtering strategies [23]. This aligns with standard practices in forensic science where different types of evidence are evaluated using the likelihood ratio method [26].

In this paper, we present a novel mathematical model that evaluates the statistical weight of pigmentation traits evidence in forensic cases. Using a likelihood-ratio approach, our model considers variables that are conditionally dependent. This is common in multiples contexts; for example, when analyzing pigmentation traits such as eye color, hair color, and skin color, assuming independence is unrealistic [27]. This aspect has not been considered by previous approaches [23]. In this work, we also measure the impact of considering these pigmentation traits in combination with genetic information when searching for missing persons. The developed methods can be applied using forensicolors, an open source R software package accessible through the CRAN repository [28].

## 2 Methods

### 2.1 Data

#### 2.1.1 Physical traits data

Physical traits, such as pigmentation, can be derived from various sources: DNA phenotyping, legal documentation, testimonies from relatives and witnesses, social media information, and direct observation. Each source possesses its own uncertainties, whether related to the methodology applied (e.g., DNA phenotyping) or to the credibility (e.g., testimonies, social media). In this context, we examine various variables collected from relatives’ testimonies in missing persons cases. For UPs, we consider DNA phenotyping or anthropological analysis. We analyze several characteristics: biological sex, denoted as *S*; age, represented as *A*; and three phenotypic variables, namely *C*_*H*_, *C*_*S*_, and *C*_*Y*_, corresponding to hair, skin, and eye color, respectively. For simplification, we introduce a composite variable, *C*, representing pigmentation traits, where *C* = *{C*_*h*_, *C*_*s*_, *C*_*Y*_ *}*. Biological sex is defined as a dichotomous categorical variable, *S* = *{F, M}*, with *F* representing female and *M* male. Age is a continuous variable ranging from 0 to 100 years. Phenotypic characteristics are polytomous categorical variables with multiple possible values. For practicality, the values are selected on the basis of the HIrisPlex classification system [4, 5, 29]. Thus, *C*_*H*_ = *{*1, 2, 3, 4*}*, with 1 for blond, 2 brown, 3 red, and 4 black hair; *C*_*S*_ = *{*1, 2, 3, 4, 5*}*, representing skin tones from very pale to dark-to-black; *C*_*Y*_ = *{*1, 2, 3*}*, with 1 for blue, 2 intermediate, and 3 brown eyes. When analyzing multiple cases, non-genetic data are stored in databases. A set of UPs is defined as *UPs* = *{UP*_1_, *UP*_2_, *UP*_3_,…, *UP*_*N*_ *}*, where *N* is the total number of unidentified persons. Each UP has associated data. Similarly, a set for MPs is defined as *MPs* = *{MP*_1_, *MP*_2_, *MP*_3_,…, *MP*_*K*_*}*, where *K* represents the total number of missing persons. For each MP, we have MP data. The gathered non-genetic information for each MP or UP constitutes the individual’s phenotype.

#### 2.1.2 Genetic data

In MPI and DVI cases, there are normally two genetic databases: (i) one for the MP relatives, named the reference genetic database, and (ii) another for the UPs, named the UPs genetic database. For each *MP*_*j*_, the family relationship is described in a pedigree *Ped*_*j*_. For each UP and MP relative, different genetic markers could be analyzed, for example, sets of short tandem repeats (STR) or autosomal single nucleotide polymorphism (SNPs) genotyped from samples (blood, bones, etc.) [30]. In this example, we consider a set of 23 STR markers for STRs. The complete genetic information obtained constitutes the individual’s genotype, *G*. Therefore, the UP genetic database has *G*_*i*_ = *{M*_1_, *M*_2_, *M*_3_,…, *M*_23_*}*, where *i* is a specific UP, and *M* are the genotypes obtained from different markers. In the same way, in the reference genetic database, *G*_*jr*_ = *{M*_1_, *M*_2_, *M*_3_,…, *M*_23_*}*, where is *j* a specific MP, and *r* relatives located in the family pedigree of *MP*_*j*_. When more genetic data are required for exceptional cases, samples could be genotyped with more autosomal STR markers, SNP, X-chromosomal STR, Y-STR, and mtDNA [2, 10].

### 2.2 Statistical evaluation of the evidence

This section presents a mathematical framework for evaluating forensic evidence. Each model contains parameters that need to be estimated or specified and are discussed in Sections 2.2.1 to 2.2.6.

#### 2.2.1 Genetic evidence

For a given UP, the kinship test involves comparing two hypotheses: *H*_1_: UP is MP, and *H*_2_: UP is not related to MP. In the MPI and DVI cases [30], this test compares, for each pair of MP-UP, the probability of genetic evidence observed under these hypotheses. This test is assessed using the LR approach, defined as:

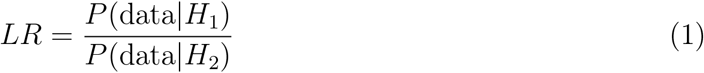

where data could be either genetic or non-genetic evidence. In a typical MPI case,the genetic profile of an individual is indicated as *G*_*α*_ = *{M*_*α*,1_, *M*_*α*,2_,…, *M*_*α,m*_*}*, where *M*_*α,i*_ is the genotype observed in marker *i* for the person *α*. Given a model defined by the marker set and parameters such as mutation rates and population structure, we can statistically evaluate the hypotheses using the LR, defined as:

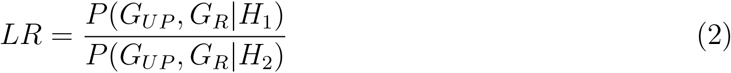

Here, *{G*_*UP*_, *G*_*R*_*}* includes genotype information for both UP and MP relatives. *P* (*G*_*UP*_, *G*_*R*_|*H*_*i*_) is the probability of observing specific genetic profiles given the scenario *H*_*i*_.

#### 2.2.2 Biological sex

In the following subsections, we introduce various LR models to evaluate non-genetic variables, such as biological sex, age, and pigmentation traits such as hair, eye, and skin colors. The models for biological sex and age have been elaborated in more detail in [23]. Also, a more general definition is available in the Appendix section. We begin with the LR computation model for the non-genetic variable, biological sex (*S*), represented as:

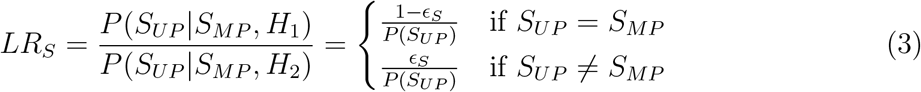

Here, *S*_*UP*_ and *S*_*MP*_ denote the biological sexes of UP and MP, respectively. The hypotheses being compared are the same as those presented for the genetic data, *H*_1_ (UP is MP) and *H*_2_ (UP is not related to MP). The model accounts for two scenarios: concordance (*S*_*UP*_ = *S*_*MP*_) and discordance (*S*_*UP*_ ≠ *S*_*MP*_). The error parameter, *ϵ*_*S*_, accounts for potential errors due to incorrect testimonies, data entry inaccuracies, laboratory estimations or misinterpretations of the source data.

#### 2.2.3 Age

The age, denoted as *A*, is a continuous variable. It incorporates uncertainties in *A*_*MP*_ arising from inaccurate testimonies and in *A*_*UP*_ due to laboratory estimates, which could include DNA-based predictions or anthropological analyses [6]. These uncertainties lead to estimated age ranges, as detailed in [31]. The age ranges are defined as *A*_*UP*_ = *{UP*_min_, *UP*_max_*}* and *A*_*MP*_ = *{MP*_min_, *MP*_max_*}*. An overlap of these age windows indicates concordance. To quantify this, a Boolean variable *ψ* is introduced, where *ψ* = 1 means overlap, and *ψ* = 0 indicates that there is no overlap. The LR model for *A* is presented below:

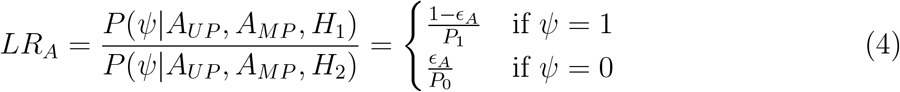

In this model, *ϵ*_*A*_ represents the error rate related to data entry errors and the uncertainties inherent in the estimation of the age range from laboratory analyzes. *P*_1_ and *P*_0_ indicate the frequency of the value of *ψ* in the reference population database. For example, if *ψ* = 1 for a specific case, *P*_1_ is the frequency of individuals in the reference population who meet the age overlap condition with *MP*. In contrast, *P*_0_ is the frequency of individuals who do not meet the overlap. Therefore, *P*_1_ + *P*_0_ = 1.

#### 2.2.4 Pigmentation traits: one variable model

Here we focus on a single pigmentation trait, hair color (*C*_*H*_). The LR es defined as:

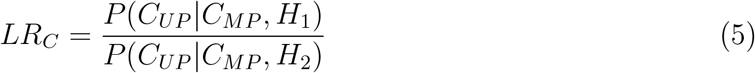

where *C*_*UP*_ and *C*_*MP*_ denote the hair colors of UP and MP, respectively. The *LR*_*C*_ can be expressed as:

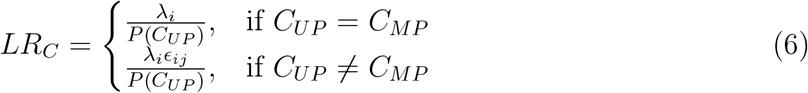

The error rate for hair color is defined for each 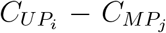 pair, reflecting the varying difficulty in hair color classification. For example, the error rate between blond and red colors (more similar) is different from the error rate between black and white colors (more different and less likely to be confused). The normalization constant *λ*_*i*_ ensures that the total probability of a given *MP* is equal to 1.

#### 2.2.5 Pigmentation traits: multiple variables model

Given a specific combination of the characteristics of the phenotype (*C*) of an MP and UP pair and defining the error rates *e*_*h*_, *e*_*s*_, and *e*_*y*_, LR can be computed as follows:

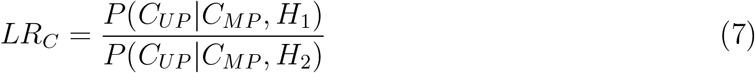

where *C* represents a specific combination of hair, skin, and eye color, *H* _1_: UP is MP, and *H*_2_: UP is not MP. The joint probability of observing a particular combination of characteristics is expressed using the chain rule of conditional probabilities.

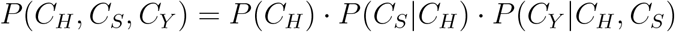

where *P* (*C*_*H*_) is the probability of observing a specific hair color, *P* (*C*_*S*_ |*C*_*H*_) is the probability of observing a specific skin color given the hair color, and *P* (*C*_*Y*_ |*C*_*H*_, *C*_*S*_) is the probability of observing a specific eye color given b oth the hair and skin colors. The LR is then defined as:

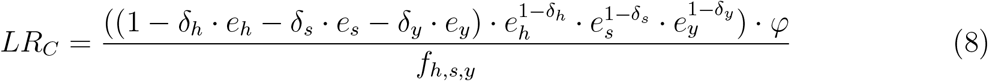

where *δ*_*H*_ is 1 if there is a concordance in hair color, and 0 otherwise; similarly, *δ*_*S*_ and *δ*_*Y*_ are 1 for correspondences in skin and eye colors, respectively, and 0 for discrepancies. The error rates for hair, skin, and eye colors are represented by *e*_*h*_, *e*_*s*_, and *e*_*y*_, respectively. *φ* is the normalization factor that corrects the numerator in order to ensure that all phenotype characteristics probabilities sum 1. The term *f*_*h,s,y*_ represents the frequency of the specific characteristic combination in the reference population, adjusted using Laplace smoothing to address unobserved attribute combinations. The frequency *f*_*h,s,y*_, is calculated by adding a count parameter *λ* to each observed frequency and to the frequencies of unobserved combinations:

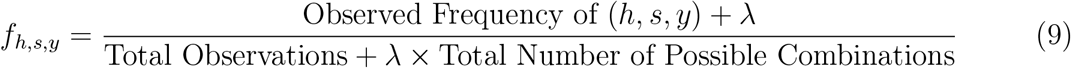

#### 2.2.6 Combining genetic and non-genetic evidence

We assume that genetic and non-genetic evidences are independent. Therefore, for a specific assignment *a*, in contrast to the null model, the combined LR is given by:

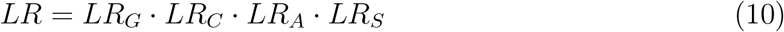

Note that this assumes that genetic, color traits, age and sex variables are conditionally independent. However, within color traits, the conditional dependency between hair, skin, and eye color is taken into account.

### 2.3 Statistical power calculation

To evaluate the statistical power of a given LR model, we define the true-positive ratio (*tp*) as *P* (*LR > T* |*H*_1_) and the false-positive ratio (*fp*) as *P* (*LR > T* |*H*_2_). Similarly, the true negative ratio (*tn*) is *P* (*LR ≤ T* |*H*_2_) and the false negative ratio (*fn*) is *P* (*LR ≤ T* |*H*_1_). These ratios are used to construct the Receiver Operating Characteristic (ROC) curve, which plots TPR against FPR at various threshold (*T*) settings.

The Area Under the ROC Curve (AUC) provides a single measure of overall performance of the LR model across all possible classification thresholds. The AUC ranges from 0 to 1, where an AUC of 0.5 denotes a model with no discriminative ability (equivalent to random chance), while an AUC of 1.0 indicates perfect discrimination between positive and negative classes.

It is important to note that these metrics were used to quantitatively describe the properties of the *P* (*LR*|*H*_1_) and *P* (*LR*|*H*_2_) distributions. In a comprehensive Bayesian framework, prior probabilities should be included, and decisions on potential correspondences would be based on posterior odds [23]. However, in this work, we focus on assessing the LR models without making assumptions about prior probabilities.

#### 2.3.1 Computational simulations

A methodology using simulation has been suggested to obtain error rates and assess performance metrics [2, 10, 19]. In this section, we describe this general approach, which is applicable to various types of evidence, both genetic and non-genetic. In the following, we present the pseudocode (Algorithm) for these simulations.

##### Algorithm Evidence simulations

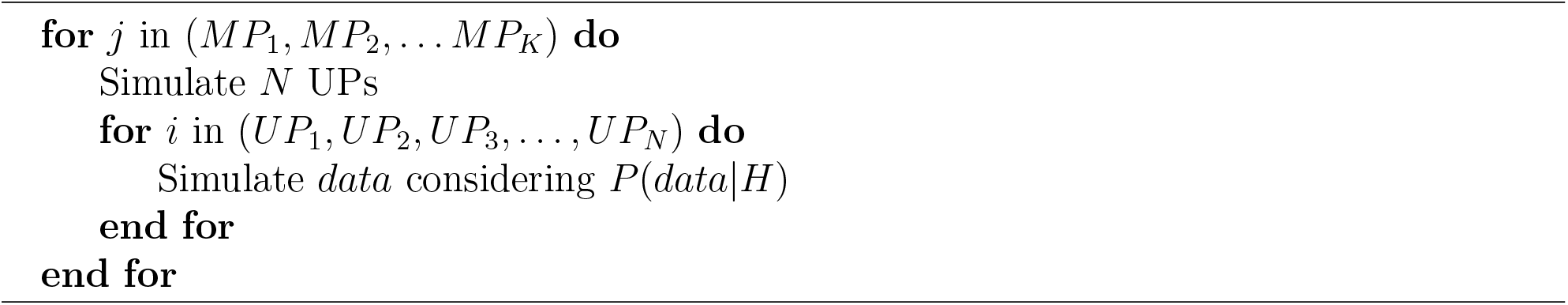

The algorithm indicates that for each *MP*_*j*_ in the database, a set of *N* UPs is simulated. Then, for each UP, *data* (that could be non-genetic or genetic information) is simulated considering *H*_1_ or *H*_2_ as true. Typically, 10,000 UPs are simulated considering *H*_1_ and 10,000 UPs considering *H*_2_. In order to present a practical example, we consider simulating the sex variable *S* for the case where *MP* is female (*S*_*MP*_ = *F*) with an error rate *ϵ*_*S*_ = 0.05. We simulate UPs resulting in two lists: (i) 10,000 UPs considering *H*_1_ as true, with expected female proportion of 0.95 (950), and (ii) 10,000 considering *H*_2_ as true, matching the population frequency *P* (*F*). These simulations could extend to other genetic and non-genetic data.

### 2.4 Implementation

All calculations presented were performed using the forensicolors R package, freely available on CRAN and developed in the context of this work. Also, `mispitools`, `forrel` R packages [2, 10], freely available in CRAN, were used to calculate genetic likelihood ratios. The handling of the pedigree was performed using the `pedtools` package [11].

## 3 Results

In this section, we present the results of applying the developed LR models.

### 3.1 Non-genetic data gathering

Different data types can be obtained from UPs (e.g., through anthropological or DNA analysis) and MPs (e.g., through testimonies, documentation, etc.). This includes the standard genetic fingerprint used in human identifications [32] and information about biological sex, age, physical characteristics, etc [1]. As shown in Figure 1a, these data are used to compare MPs with UPs, occasionally through database search [33]. Here we performed an analysis based on previously developed LR models [23] for preliminary investigation data as an introduction to the results section. Figures 1 a-c show the expected distributions of the LR values for different variables such as biological sex (Figure 1a), hair color (Figure 1b) and age (Figure 1c) according to the hypotheses *H*_1_: UP is MP and *H*_2_: UP is not MP. Generally, higher LR values are expected with *H*_1_. However, various errors such as data entry errors, methodological misclassifications in anthropological analysis, and uncertainty in testimony, among others, can result in low LR values even when *H*_1_ is true. This is exemplified in Figure 1B, where a small number of cases could produce LR = 0.1 under *H*_1_. Ignoring potential error types in the LR model can lead to misleading conclusions, as discussed in [34]. If the different pieces of evidence are conditionally independent, a combined LR can be computed by multiplying the individual LR, as illustrated in Figure 1c. This improves the distinction between the expected LR values under *H*_1_ and *H*_2_, improving the discrimination capacity based on LR [2]. We next turn our attention to pigmentation traits, applying the proposed model to statistically weigh the evidence while considering trait interdependence.

**Figure 1:**
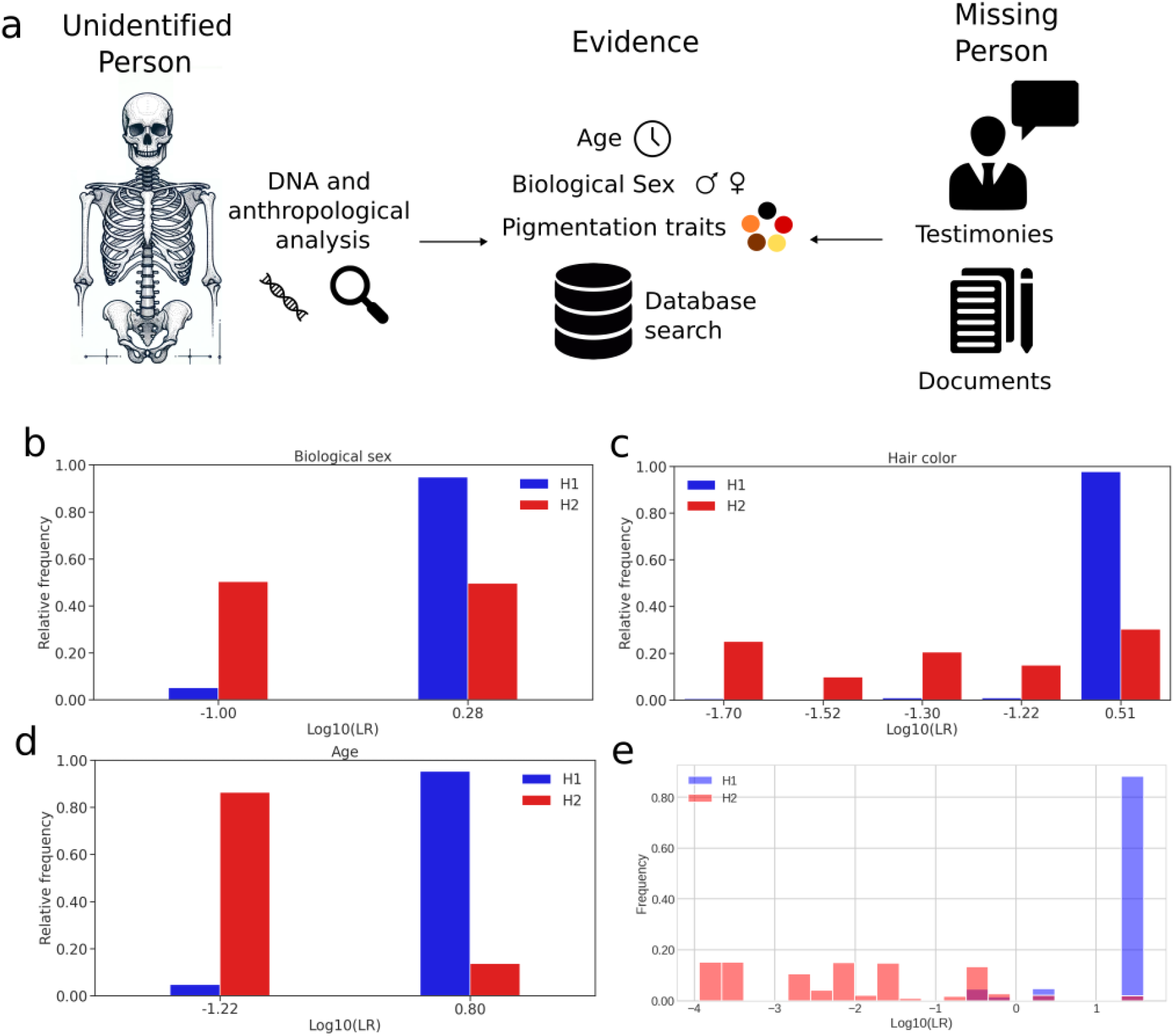
a Evidence gathering during a missing person or disaster victim identification case. Some data are collected from the unidentified remains and interpreted through DNA or anthropological analyses leading to age, biological sex and pigmentation traits estimation between others. Also, the same data are collected from the missing person, generally from relatives testimonies and documentation. All data are stored and compared through databases. b - Likelihood ratio distribution for biological sex variable. c - Likelihood ratio distribution for hair color. d - Likelihood ratio distribution for age. e - Combined Likelihood Ratio distribution considering all previous variables. In all cases, expected values considering *H*_1_ are in blue, and those expected considering *H*_2_ are in red. In all models, error rate, *ϵ* = 0.05 was used.

### 3.2 Likelihood Ratio model for pigmentation traits

As stated in Equation 1, the LR computation requires defining the conditioned probabilities of observing the data under each hypothesis. In Figure 2, the conditioned probabilities are illustrated. Figures 2a and b show *P* (*D*|*H*_1_). Thus, these probabilities depend on the characteristics of a specific MP. We introduce two MPs: *MP*_1_ with eye color 2, skin color 3, and hair color 2; and *MP*_2_ with eye color 3, skin color 5 and hair color 3. In all cases, the error rate *ϵ* = 0.01 was used (see Equation 8). When searching for conditioned probabilities in both cases, the maximum likelihood is obtained with UPs possessing identical characteristics (0.84 in both cases). Furthermore, there is a probability of 0.017 for several possibilities associated with just one discrepancy in the variables (e.g., different hair color but same eye and skin color). A lower probability (0.00035) is observed for cases with two discrepancies and the lowest probability (0.0000071) for those with three discrepancies, indicating a small but non-zero probability even when no characteristics have the same value. Lastly, for *P* (*D*|*H*_2_): UP is not MP; applying Equation 9 to a simulated traits frequency database (available in Supplementary Materials) yields the frequencies shown in Figure 2c. This shows that some characteristics are more prevalent than others, notably, the combination of characteristics of *MP*_2_ is particularly rare in this example.

**Figure 2:**
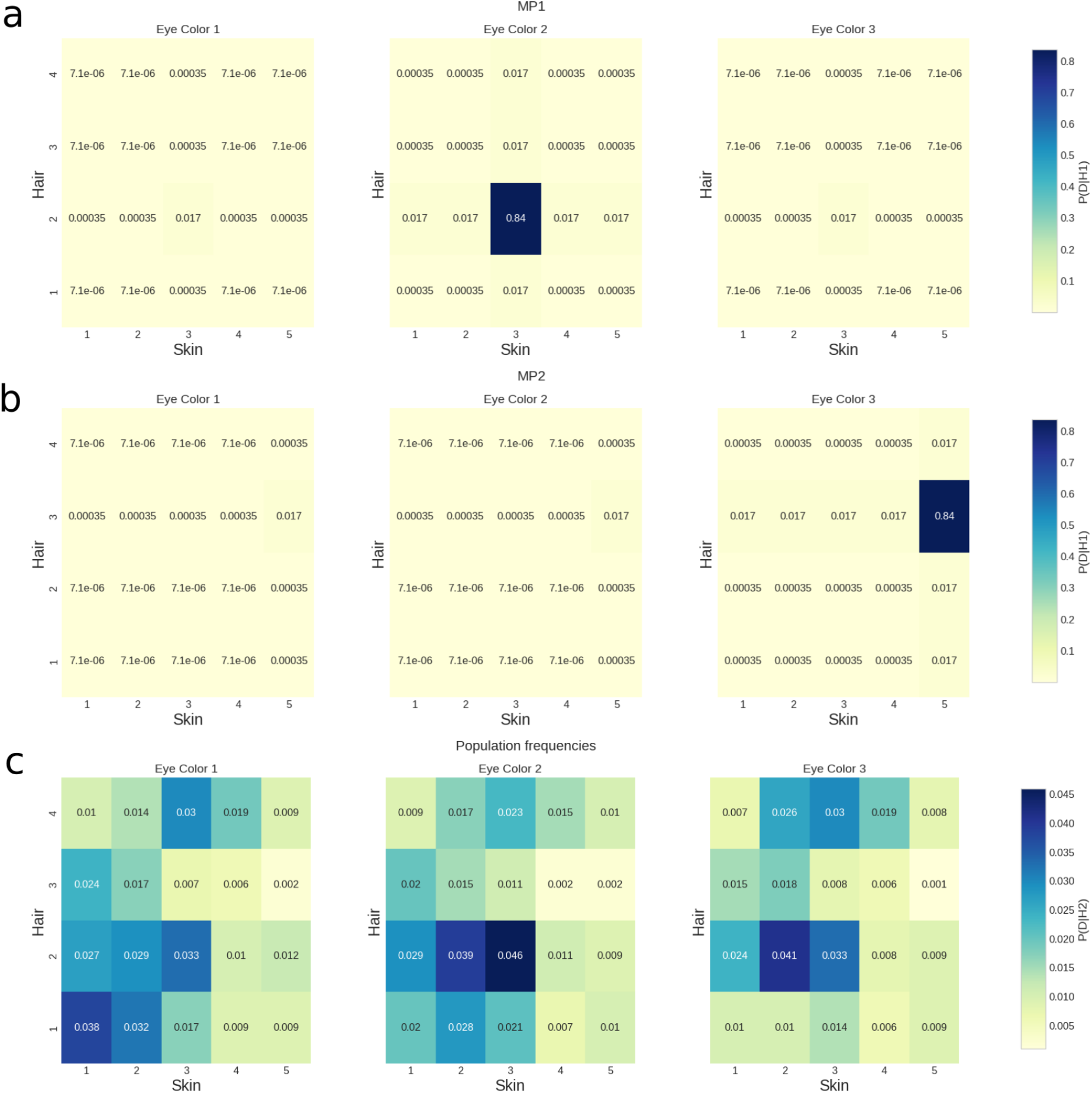
a - Conditioned probability table for pigmentation traits considering *H*_1_ as true and *MP*_1_. This probability distribution is obtained from the numerator of Equation 8, considering the characteristics of *MP*_1_. It shows the probability of finding UPs with the specified characteristics given that UP is *MP*_1_ (*H*_1_ as true). b - The same conditioned probability table considering *MP*_2_ characteristics. c - Conditioned probability table for pigmentation traits considering *H*_2_ as true. This probability distribution is obtained using equation 9. Briefly, it shows the probability of finding individuals with the specified characteristics in the reference population.

Once the likelihoods are defined, the LR can be computed for each characteristic using Equation 8. For example, with *MP*_2_, analyzing a case where *UP* shares the same characteristics, 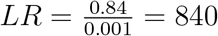 is obtained. In addition, conditional probability tables can be utilized to simulate possible pigmentation traits under given hypotheses. For example, considering *H*_1_ as true (Figure 2b), it is expected that in 84% of the cases, *LR* = 840 will be obtained, *P* (*LR* = 840|*H*_1_) = 0.84. In contrast, if *H*_2_ is true (Figure 2c), only 0.1% are expected to report a *LR* = 840, *P* (*LR* = 840|*H*_2_) = 0.001. This is an expected behavior, since *H*_1_ states that UP is MP, therefore higher LR values (i.e. 840) are more likely to be obtained compared to the case where *H*_2_ (UP is not MP) is considered true. This method enables the generation of LR distributions under different hypotheses, as shown in Figures 3a and 3b. Furthermore, Receiver Operating Characteristic (ROC) curves can be derived by considering different thresholds. For both MPs, high Area Under the Curve (AUC) values are observed when considering only pigmentation traits, as shown in Figure 3c.

**Figure 3:**
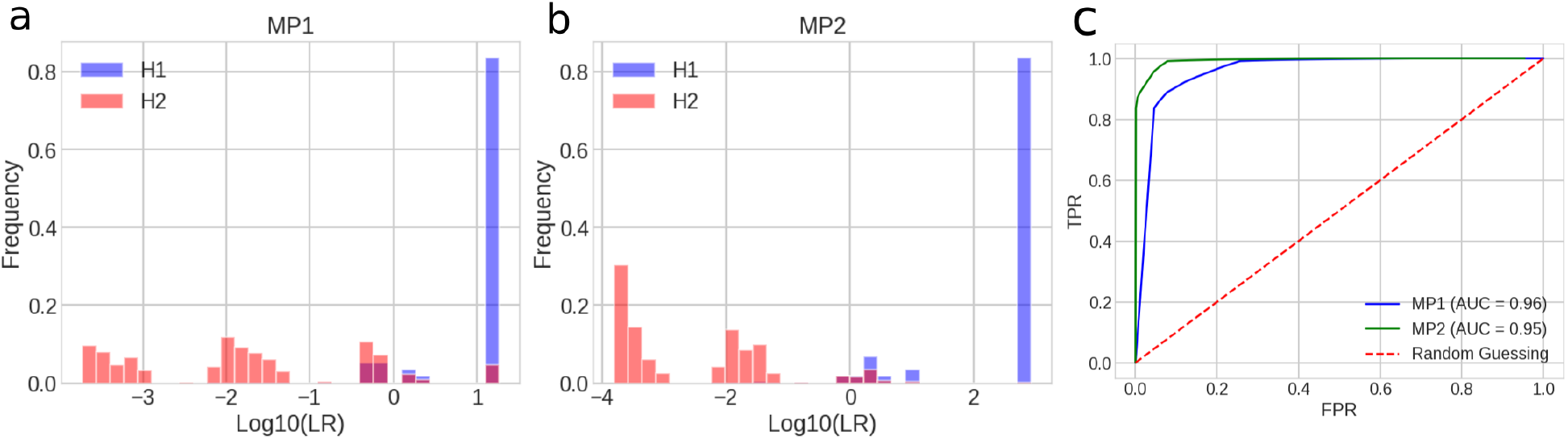
a - *Log*_10_(*LR*_*C*_) distributions considering *H*_1_ and *H*_2_ as true for *MP*_1_. b - *Log*_10_(*LR*_*C*_) distributions considering *H*_1_ and *H*_2_ as true for *MP*_2_. In both cases, *H*_1_ expected values are in blue and *H*_2_ are in red. c - ROC curves for both distributions, in red the random classifier is shown.

### 3.3 Combining DNA with pigmentation evidence: application in MPI cases

In the previous section, relying solely on non-genetic (or genetically predicted) pigmentation traits yielded acceptable performance. In fact, forensic searches sometimes begin with only this type of information [1]. However, in scenarios where only a few or distant relatives of the missing person are available, standard forensic 23 autosomal STRs may not provide sufficient statistical power for search [10, 2]. In such cases, other sources of evidence become crucial [23]. We analyze a group of under-powered pedigrees and combine DNA-based LR (*LR*_*G*_) with those derived from pigmentation traits (*LR*_*C*_) (Figure 4). It was observed that the use of genetic information only results in overlapping LR distributions under *H*_1_ and *H*_2_ (Figure 4). This indicates the potential for high LR when UP is not MP and low *LR*_*G*_ when UP is MP, leading to probable false positives and negatives. The most pronounced example is shown in Figure 4c, where the distributions *LR*_*G*_ almost completely overlap with only a second cousin. In every scenario, as expected, combining *LR*_*C*_ with *LR*_*G*_ (for both *MP*_1_ and *MP*_2_) leads to a separation of distributions, reducing the probability of high LR values when UP is not MP (potential false positives) and low values when UP is MP (potential false negatives).

**Figure 4:**
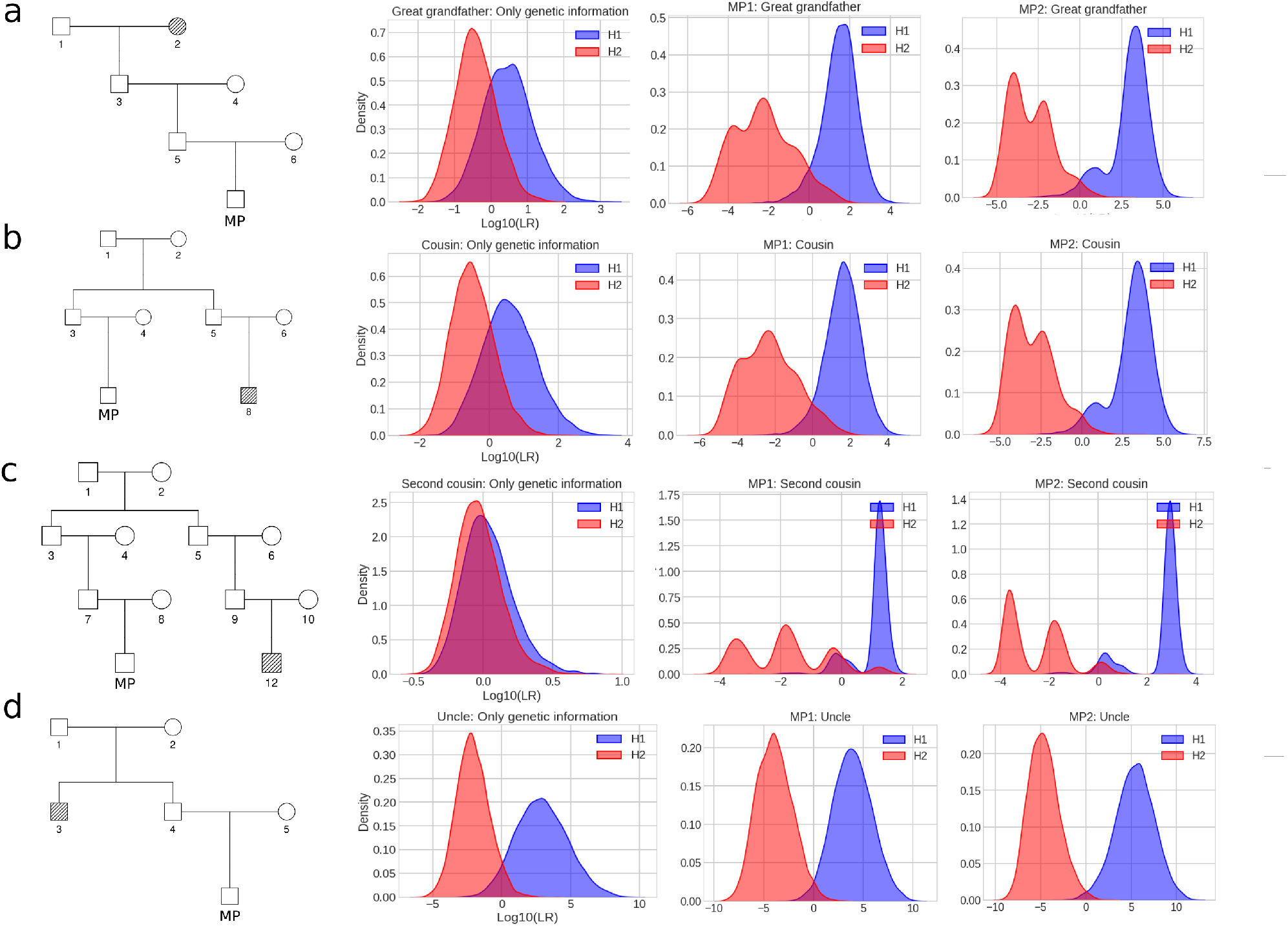
a - Pedigree, *LR*_*G*_, combined *LR* for *MP*_1_ and combined *LR* considering *MP*_2_ are shown for a pedigree with a great grand-father genotyped. b - The same distributions but for a pedigree considering a typed cousin. c - In this case, only a second-cousin is genotyped. d - A case where the uncle is genotyped. In all cases missing individuals are in red.

This effect is also reflected in the ROC analysis shown in Figure 5. In all cases, the ROC curves show better performance for the combined *LR*. This could be analyzed through the AUC (Table 1). Higher AUC is obtained always when the phenotype data is incorporated. In particular, *MP*_2_ shows better performance, which could be attributed to its characteristics of the less common pigmentation traits.

**Figure 5:**
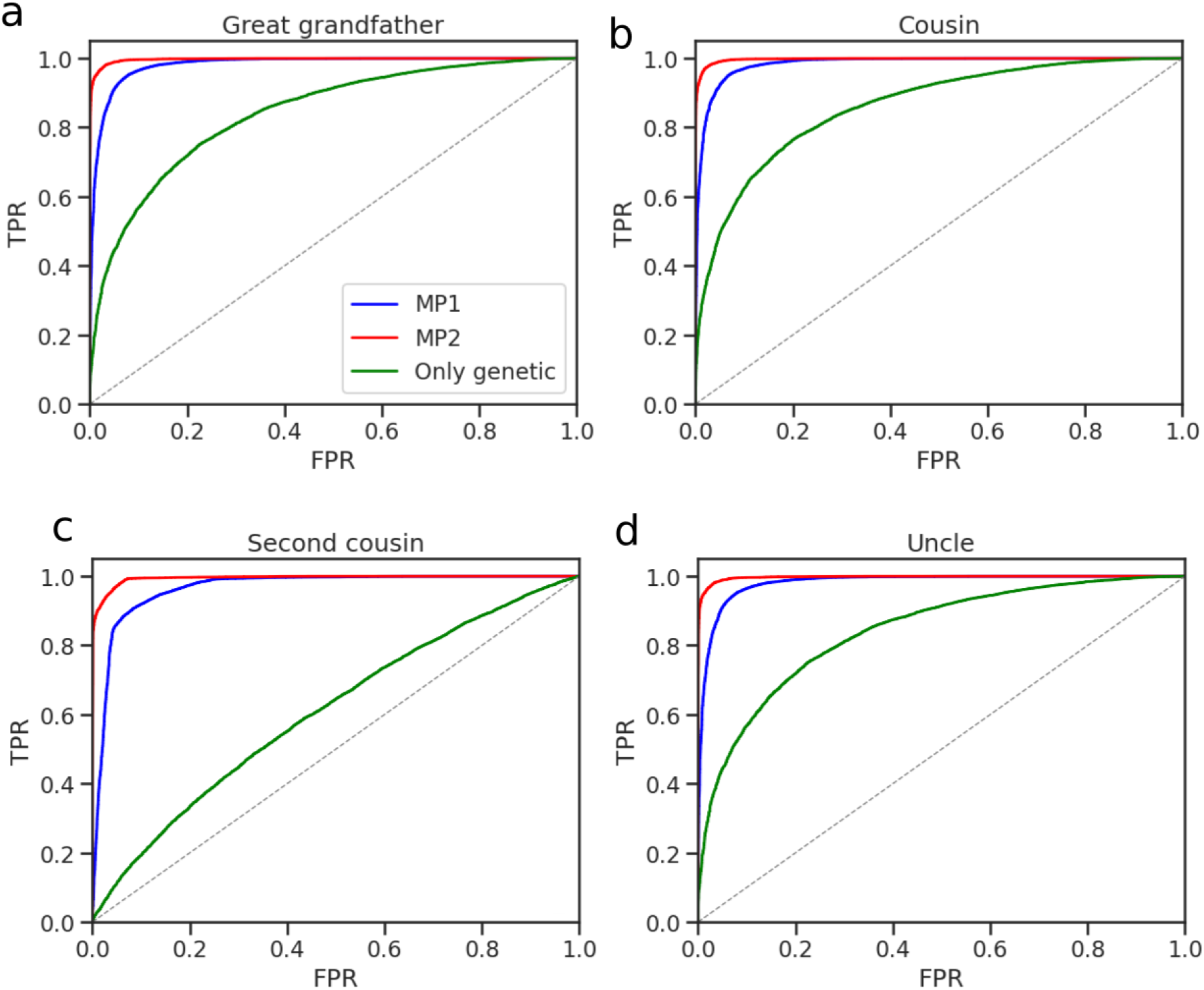
ROC curves from pedigrees analyzed in Figure 4. a - Great grandfather, b - Cousin, c - Second cousin, d - Uncle. Dashed line represent a random classifier. Blue curve is for *LR*_*G*_, red curve for combined *LR* considering *MP*_1_ and green curve for combined *LR* considering *MP*_2_.

**Table 1:**
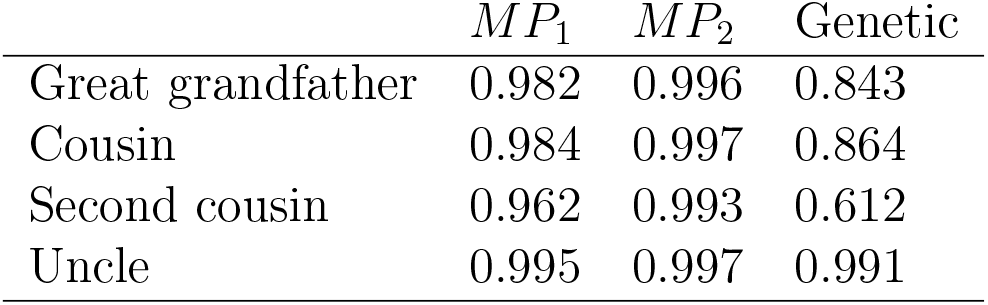
Area Under the Curve obtained from ROC plots present in Figure 5. Genetic refers to the AUC obtained from the green curve, *LR*_*G*_. *MP*_1_ from red curve, combined *LR* considering *MP*_1_, and *MP*_2_ refers to the green curve, combined *LR* considering *MP*_2_.

## 4 Discussion

In this study, we have developed mathematical models that enable the computation of a statistical weight of evidence for a comprehensive range of non-genetic variables. In particular, we focus on the characteristics of the pigmentation traits. Due to recent advances in DNA phenotyping [3, 4, 6, 29] and the widespread adoption of digital devices that facilitate the acquisition of individual photographs, among other types of information [35], the application of pigmentation traits characteristics in the search process for missing people is on the rise [1]. Despite this, there is a notable lack of mathematical models for computing likelihood ratios. As described by Evett and Weir [8, 36], a fundamental role of the forensic scientist is to provide a statistical weight of the evidence. This formalization, as previously demonstrated in [23], allows the incorporation of various phenomena such as typing errors and methodological inaccuracies, which, if not properly accounted for, could lead to misleading conclusions. Furthermore, an appropriate formalization of the statistical weight of evidence could serve as an initial step towards integrating different types of evidence, which could significantly improve forensic tasks [37].

Previous works have discussed the use of non-genetic data in missing person searches [14, 20, 21, 38]. Specifically, Budowle et al. [14] underscored the need to develop guidelines for objectively computing prior odds based on non-genetic data, such as phenotypic characteristics. In their study, the authors reviewed various approaches for setting prior odds, highlighting the risks associated with potential data misuse. They advocated for a consensus on accounting for the uncertainty inherent in eyewitness accounts. However, Biederman et al. [39] argued that establishing guidelines or an objective approach to prior odds is impractical, since prior probabilities are inherently expressions of personal and thus subjective beliefs. Our proposed methodology aims to address this issue by treating non-genetic characteristics as evidence. We suggest models for measuring the statistical weight of the evidence by likelihood ratios, in a similar way as the methods used for DNA data.

Our methodology addresses pigmentation traits, demonstrating the benefits of this approach in terms of reliability and consistency. Our model distinguishes itself from previous ones that worked with non-genetic variables in these contexts [14, 20, 21, 23] by incorporating conditional dependencies between variables, extending the framework to a more realistic modeling.

The presented methodology is particularly effective in scenarios where genetic evidence is sparse or nonexistent. It allows for the strategic selection of cases for further investigation, gathering either non-genetic or genetic data. This approach is highly applicable in missing person searches driven by non-genetic data from missing persons (*MP*), a concept demonstrated in our results section. On the other hand, our methodology can also be adapted to prioritize based on data from an unidentified person (*UP*). This prioritization is essential for the identification of disaster victims, where a particular remains may need to be compared with multiple relatives *MP*. In situations with limited genetic data from *MP* relatives, our approach highlights the importance of using non-genetic data to identify cases where additional genetic analysis is most urgently needed. Moreover, DNA-based solutions can exhibit symmetry. For example, in scenarios where two siblings are missing and only parental genetic data is accessible, DNA analysis alone may be insufficient to ascertain the identity of each missing individual. In such cases, using non-genetic data, such as age, can resolve these cases, occasionally with certainty. If age estimation carries inherent uncertainty, LR can also be computed to reflect non-genetic evidence [23].

However, there are several limitations to consider when applying the models we have presented. The improvement in adding information on pigmentation traits depends heavily on the reference population database. One primary concern is the frequencies of the reference population for phenotypic characteristics. Current research aims to define pigmentation traits more accurately [40, 41]. We anticipate that advancements in the computerization and automation of missing person search databases will yield more reliable and accurate measurements for the frequency of these characteristics. Moreover, we know that the categories selected for defining pigmentation traits are just a reduction of the diverse color palette that represent the possible tones that describe the human diversity [42]. In future works this diversity should be incorporated. Another critical aspect is that our model assumes a generic error rate that can be easily adjusted for various technologies and data sources. Moreover, calibrating different DNA phenotyping approaches could lead to a more precise understanding of the accuracy of the methodology, enabling a more informed estimation of error rates. Nevertheless, knowing the error probability from the testimony of a missing person’s relative may not be possible in many cases. We encourage forensic professionals to adopt a conservative approach in such cases, setting error rates that avoid over reliance on the source and thus not discounting the possibility of errors, as previously discussed in [23]. An important point is that adding phenotype information does not replace the DNA-based framework; moreover, it could be used as a complementary tool. This strategy could be combined with other approaches to lead with low statistical power in large-scale cases [2, 10, 43].

Recently, advances in technology have significantly improved phenotyping capabilities [6], facilitated the extraction of data from digital sources [35], and improved voice recognition techniques [26]. These innovations play a crucial role in forensic science, where experts in various disciplines are dedicating efforts to statistically evaluate diverse evidence sources [26]. The integration of evidence from different sources, viewed through the lens of mathematical formalization, is a vital step forward. This approach is not only about combining data but also about enhancing the reliability and accuracy of the information used in the search process. Using advanced statistical models and analytical tools, we can synthesize information from various fields, such as forensic entomology [44], crime scene investigations[45], voice analysis [46], and traditional investigative methods. This comprehensive methodology aims to improve the effectiveness of searches for missing persons, disaster victim identification, and criminal investigations, particularly in complex and challenging scenarios. The goal is to establish a more robust framework that can adapt to the dynamic nature of forensic investigations [1].

### 5 Conclusion

This paper proposes a model for calculating likelihood ratios using a set of phenotypic variables, particularly pigmentation traits. Moreover, we have shown the impact of incorporating these likelihood ratios in database search performance through computational simulations. The calculation algorithm and functions are implemented in `forensicolors`, freely available on CRAN.

## 6 Appendix: General likelihood

Consider the non-genetic evidence. We have observed a feature vector in UP and MP, denoted

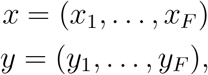

respectively. We consider the likelihoods *L*(*x* | *y, H*_1_) and *L*(*x* | *y, H*_2_). The latter simplifies to *L*(*x*) and therefore we only need to specify or estimate the relative frequency of *x* in the relevant population. Assume first that features are conditionally independent and consider

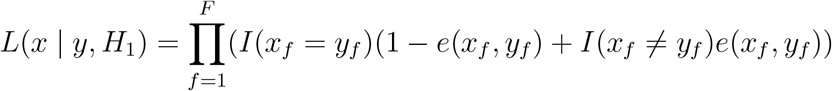

where *I*() is the indicator function and *e*() is the error function. In the simplest case, similar to several previous examples, *e*(*x*_*f*_, *y*_*f*_) = *ϵ*_*f*_ if *x*_*f*_ = *y*_*f*_ and 1 *− ϵ*_*f*_ otherwise. We could, however, use more refined versions to allow for some errors to be less likely than others. For example, *e*(*x*_*f*_, *y*_*f*_) could depend on the distance between the feature values, i.e. | *x*_*f*_ *− y*_*f*_ |, for features with more than two levels. The above assumption of independence can also be relaxed. We would then replace *e*(*x*_*f*_, *y*_*f*_) by *e*(*x*_*f*_, *y*_*f*_ | *A*_*f*_) to indicate conditioning on the characteristics preceding f-th.

## Notes

https://github.com/MarsicoFL/forensicolors

### Competing Interest Statement

The authors have declared no competing interest.

