## Supplementary Material for "Likelihood Ratios for physical traits in forensic investigations"

### Forensicolors

Tutorial

13th May 2024

In this tutorial we introduce basic commands provided by the R package forensicolors. This package Computes likelihood ratios based on pigmentation traits. Also, it allows computing conditional probabilities for unidentified individuals based on missing person characteristics. Tailored plots are incorporated to analyze likelihood ratio distributions.

#### Installation

Firstly, R (≥ 2.10) should be previously installed. It is possible to execute forensicolors using the R command line or also Rstudio (https://posit.co/download/rstudio-desktop/). One opened, type the following commands in the R console:

```
> install.packages("forensicolors")
> library(forensicolors)
```

This will install and load the forensicolors package. Also other dependencies (if they are not previously installed) will be downloaded. Particularly, forensicolors depends on the following R packages: forrel, pedtools, plotly, dplyr, ggplot2.

#### Obtain the reference population data for pigmentation traits

The first step in a basic analysis is to simulate a reference population database. It is done through the function simRef(). It could be executed as following:

```
> data <- simRef()
```

This will use the default parameter values, for more information about the function:

```
> ?simRef()
```

#### Compute UPs conditional probabilities based on MP characteristics

Then, the conditional probabilities for UPs based on MPs characteristics (given H1) could be computed using contionedProp() function. This allows users to enter as parameters different values for hair, skin and eye colors. Also, it allows the introduction of error rates. The function could be executed as following:

```
> conditioned <- conditionedProp(data, 1, 1, 1, 0.01, 0.01, 0.01)
```

In this example, hair, skin and eye colors values are 1. For more details ?conditionedProp() could be executed.

<u>Compute pigmentation traits probability distributions based on reference population data</u>

Once simulated the reference population, the probability distributions based on reference population data (given H2), could be computed as following:

```
> unconditioned <- forensicolors::refProp(data)
```

Note that the parameter *data* comes from the command previously executed above (> data <- simRef()).

<u>Compute LRs</u>

With both probability distributions previously calculated, we can now compute the Likelihood Ratios using the following command:

```
> likelihoodsR <- compute_LRs(conditioned, unconditioned)
```

<u>Plot LR distributions considering both H1 and H2 as true</u>

Finally, a specific function for plotting the likelihood ratios is defined below:

```
> plotLR(likelihoodsR)
```
